# Including Data Analytical Stability in Cluster-based Inference

**DOI:** 10.1101/844860

**Authors:** Sanne P. Roels, Tom Loeys, Beatrijs Moerkerke

**Affiliations:** Department of Data Analysis, Ghent University, Henri Dunantlaan 2, 9000 Ghent, Belgium

**Keywords:** fMRI, testing, stability

## Abstract

In the statistical analysis of functional Magnetic Resonance Imaging (fMRI) brain data it remains a challenge to account for simultaneously testing activation in over 100.000 volume units or voxels. A popular method that reduces the dimensionality of this test problem is cluster-based inference. We propose a new testing procedure that allows to control the family-wise error (FWE) rate at the cluster level but improves cluster-based test decisions in two ways by (1) taking into account a measure for data analytical stability and (2) allowing a more voxel-based interpretation of the results. For each voxel, we define the re-selection rate conditional on a given FWE-corrected threshold and use this rate, which is a measure of stability, into the selection process. In our procedure, we set a more liberal and a more conservative FWE controlling threshold. Clusters that survive the liberal but not the conservative threshold are retained if sufficient evidence for voxelwise stability is available. Cluster that survive the conservative threshold are retained anyhow, and clusters that do not survive the liberal threshold are not further considered. Using the Human Connectome Project Data (Van Essen et al., 2012), we demonstrate how in a group analysis our method results not only in a higher number of selected voxels but also in a larger overlap between different test images. Additionally, we demonstrate the ability of our procedure to control the FWE, also in relatively small sample sizes.

## 1 Introduction

Following every scientific experiment the researcher is entrusted with weighing the cumulated evidence and inferring correct and relevant information. For Functional Magnetic Resonance Imaging (fMRI) data, signals are measured via the Blood Oxygen Dependant (BOLD) signal. Evidence for brain activation is typically summarized in a statistical parametrical map (SPM) or a test image based on the general linear model (GLM) (e.g. Lindquist, 2008). These images consist of a summary statistic for each of the > 100.000 voxels, i.e. the small volumetric units that form the brain volume, and can be a summary of a single subject study, a group study (multi-subject study) or a meta-analysis. In each voxel, the evidence against the null hypothesis of no activation (*H*_0_) is tested. When *H*_0_ is rejected, this provides evidence for the alternative hypothesis of true activation (*H*_1_).

As the amount of voxels is large, correcting for the multiplicity of tests is necessary to control the number of false positives (FP, see also Table 1). While voxel-based corrections can result in overtly conservative results (Worsley, 2007; Nichols and Hayasaka, 2003), cluster-based inference is an elegant approach to reduce the dimensionality of the test problem (e.g. Forman et al., 1995). Despite concerns on interpretation of such clusters and the statistical validity of the test results (Eklund et al., 2016; Woo et al., 2014; Hayasaka and Nichols, 2003), Woo et al. (2014) showed the continuing popularity of the method.

**Table 1.** Table of events for Null Hypothesis Significance Testing (NHST) in which evidence against a null hypothesis *H*_0_ is evaluated in the direction of an alternative hypothesis *H*_1_.

|  |  | Decision |  |
| --- | --- | --- | --- |
| | | Conclude $H_0$ | Conclude $H_1$ |
| Truth | Active | False Negative (FN) Type II error | True Positive (TP) |
|  | Inactive | True Negative (TN) | False Positive (FP) Type I error |

In cluster-based inference, the feature of interest is a cluster, which is defined as a collection of neighboring voxels that survive a first threshold *T*_*u*1_ (cluster-forming threshold) on a test image (Brett et al., 2007). To determine the probability to observe a given cluster with size *S* under the *H*_0_ of no activation, given this first threshold, one typically relies on a fast Random Field Theory based approximation (RFT; Friston et al., 1996). This RFT approach relies on several assumptions that are in practice very hard to verify, especially with regard to the smoothness, and the height of *T*_*u*1_ (Hayasaka and Nichols, 2003; Eklund et al., 2016, 2018).

In cluster-based testing, over 100 clusters are typically evaluated simultaneously. Two corrections for multiple testing are dominant in literature, Family-Wise error rate (FWE) control (e.g. Brett et al., 2007) and False-Discovery Rate (FDR) control (e.g. Chumbley and Friston, 2009; Chumbley et al., 2010). FWE control uses the null distribution of the maximum cluster size *max*(*S*) to control the probability of *at least* one false positive. The FDR control uses the expected fraction of false positive clusters over all findings (i.e. TP and FP) and relies on the observed *p*-values per cluster to control the nubmer of FP (Benjamini and Hochberg, 1995, e.g.).

Small sample sizes and the huge dimensionality of the test problem result in low statistical power (Carp, 2012; Button et al., 2013), making studies prone to miss activation (see e.g. Button et al., 2013; Durnez et al., 2014b). If one is however prepared to allow for more FP’s, this will increase statistical power as the number of FN’s will become smaller. By the dimensionality of the multiple testing correction problem the interplay between FP and FN is more complicated compared to a standard statistical testing situation (Mumford, 2012; Durnez et al., 2014b). Given the widespread appreciation of cluster-based inference, but the difficult balance to control errors, we propose to add voxelwise information on the stability of the results to the testing procedure. data analytical stability can be measured through the variability on the number of selected features (e.g. voxels, clusters) when the same threshold for inference is used in a replication context (Qiu et al., 2006). Results that show a higher variation, can be considered as less stable. Using the approach via testing, the concept was initially introduced in genetic association studies (e.g. Qiu et al., 2006; Gordon et al., 2007) but recently extended to fMRI (Durnez et al., 2014a; Roels et al., 2015, 2016). Roels et al. (2016) demonstrated that for group studies, voxelwise FWE and FDR corrected analyses resulted in the same ROC curve and hence on average an equal trade-off between FP and FN (see also e.g. Durnez et al., 2014b). However, FDR corrected analyses resulted in a higher variability on the number of selected voxels (see also Durnez et al., 2014a; Qiu et al., 2006). We believe that data analytical stability is an informative addition for the evaluation of test procedures, in which the current emphasis is mostly on average performance only. Furthermore, as the stability can be derived not only for the clusters, but also for the voxels, it potentially can improve the interpretation of cluster-based analysis.

While Qiu et al. (2006) coined the term stability in an inferential framework of genetic testing, Meinshausen and Buühlmann (2010) introduced stability selection more generally in high dimensional settings with highly correlated and noisy variables. The criterion is based on the determination of the selection rate over a finite number of bootstrap samples from the original sample. Consider the following general strategy. To a select a stable set of features from a data set with a given selection mechanism, (e.g. the selection of a predictor *x*_*p*_ from a set of *p >> n* highly correlated variables), set up *K* bootstrap samples and consider the set of selected features for each bootstrap sample with fixed parameters for the selection algorithm. Determine over the *K* bootstrap samples how many times all features are selected and retain only the features exceeding a selection probability above a specific cut-off *π*_*thr*_, with 0 < *π*_*thr*_ < 1. Meinshausen and Buühlmann (2010) further demonstrated that rational choices for *π*_*thr*_ (e.g. 0.6 < *π*_*thr*_ < 0.9) do not impact results substantially.

Following this rationale, it is useful to consider the construction of voxel-wise re-selection rates (Roels et al., 2016) to select stable activation regions. These rates allow to quantify the stability of a voxel, given a fixed thresholding method, and have previously been added to the inference procedure within genetic association studies (Gordon et al., 2009). Furthermore, as these rates can be computed for every thresholding method, they have the potential to add useful voxelwise information to cluster-based inference. Indeed, one major restriction of cluster-based inference is the lack of a voxel-based interpretation of the results (Nichols, 2012; Woo et al., 2014; Durnez et al., 2014b). As a significant cluster can only be interpreted in such a way that *in at least one voxel, somewhere in the cluster, there is non-zero signal* (Nichols, 2012; Poldrack et al., 2011), the procedure becomes less attractive. By adding information on the data analytical stability of a cluster, this can be partially resolved. As results of cluster-based inference have been shown to be unstable with low degrees of smoothness (Hayasaka and Nichols, 2003; Nichols, 2012), information of the stability may be advantageous in these situations. Quantification of this (in)stability and adding this into the decision process may further improve cluster-based inference.

Here, we propose a procedure that aims to control the amount of FP in a principled way, but also allows to increase ability to detect activation. In general, we determine 2 principled thresholds for cluster size at different levels *α*. As FWE correction was found to be less variable than FDR corrections (Qiu et al., 2006; Durnez et al., 2014b; Roels et al., 2016), we determine two FWE controlling thresholds for cluster size at cluster level: 1) a first relatively conservative threshold and 2) a more liberal threshold. Clusters above the conservative threshold are considered as strong evidence for activation and are retained. In a last step, we use the stability in the selection process for activation. Clusters with a *p*-value lower than the liberal threshold, but higher than the conservative threshold are retained if in addition high stability is demonstrated.

In section *Method* we describe the proposed method in depth, in *Evaluation of the procedure* we evaluate our procedure using data from the Human Connectome Project (Van Essen et al., 2012) for two purposes. First we focus on the FWE control for FP using data without systematic activation, and next we focus on activation data to derive the operating characteristics for the method to detect activation. Then we present the results and we conclude with a discussion.

## 2 Method

In this section, we first briefly describe the mass-univariate GLM approach to analyse fMRI data which is the foundation for cluster based inference. Next, we propose our new method that incorporates cluster size in a formal way via deriving minimum cluster sizes based on RFT inference. At last, the voxel-wise data analytical stability is considered.

### 2.1 Mass-univariate GLM

In a first stage, a GLM is fitted per subject (no index for the ease of notation) for the BOLD signal of each voxel over time **y**_**v**_ (**y**_**v**_ : *y*_*v*1_, … *y*_*vn*_, … *y*_*vN*_) with *N* : total number of time points, and with *v* = 1, …, *V* the total number of voxels in the brain volume (see e.g. Kiebel and Holmes, 2007; Worsley et al., 2002; Poline and Brett, 2012).

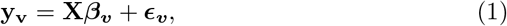

In Equation (1) **X** is a matrix that represents the expected BOLD signal under brain activation. This is a convolution of the stimulus onset function(s) with the hemodynamic response function (HRF) (Henson and Friston, 2007) for the BOLD signal. ***∈***_***v***_ is the vector representing the residuals per voxel *v*. The estimands of interest are usually single parameters of the ***β***_***v***_ vector or a linear contrast of several parameters within ***β***_***v***_. These quantities are typically estimated based on weighted least squares estimation procedures that account for the temporal dependency (Cochrane and Orcutt, 1949; Kiebel and Holmes, 2007; Worsley et al., 2002).

In a single subject analysis, a *T* map or SPM is obtained based on these estimators and standard errors for each voxel. For group or multi-subject analyses, these estimators are transferred to the group level. Consider for each voxel an estimator 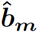 (*m* : 1 … *M* with *M* the number of subjects) as the input contrasts for the group level (Beckmann et al., 2003). A GLM is used to weight the evidence over the *M* subjects (e.g. Mumford and Nichols, 2006):

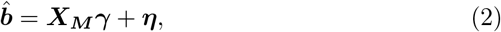

where ***X***_***M***_ denotes the design matrix. After the estimation of *γ* (Beckmann et al., 2003; Mumford and Nichols, 2009; Worsley et al., 2002), a test statistic *T* is computed for each voxel.

### 2.2 Cluster-based inference including data analytical stability at voxel level

Our proposed method to select clusters sets two thresholds and incorporates the average re-selection rate of the cluster. We stipulate how our principled procedure allows for a *deliberation* of clusters that only survive the liberal threshold by the inclusion of the stability in the test procedure.

#### 2.2.1 Thresholds based on random field theory

We define the thresholds using Random Field Theory (RFT) approximations for FWE corrected *p*-values (Worsley, 2007). RFT conveniently allows to approximate the distribution of the extent of a cluster *S* as well as the distribution of the maximum extent *max*(*S*).

A cluster is defined as a collection of neighbouring voxels that exceeds a first threshold *T*_*u*1_. The cluster extent in a Gaussian random field with dimension *D* can be re-formulated as *S ≈ cH*^*D*/2^ (Worsley, 2007) where H denotes the quadratic of height above threshold *T*_*u*1_ and where *c* equals:

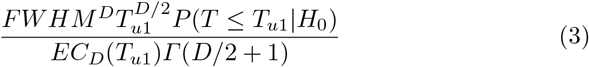

with *EC*_*d*_(*t*) the *d*-dimensional Euler Characteristic density of the test statistic *t* and *Γ* the gamma function. The Full-Width Half Maximum (*FWHM*) describes the width of the smoothing kernel that should be applied on a dataset to achieve the same amount of smoothing in data.

The probability to obtain a cluster of size *S* under the *H*_0_ of no activation for a given first threshold *T*_*u*1_ and a given *FWHM* is approximated by

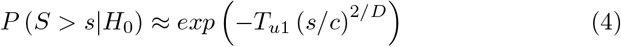

Based on these approximations, it is possible to derive FWE-corrected *p*-values.

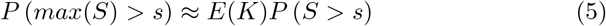

with

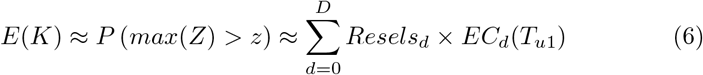

with *Resels*_*d*_ the number of *d*-dimensional or resolution elements (Brett et al., 2007, p. 232) which can be estimated from the data. To select clusters based on the *p*-values derived in Equation (5), one defines a threshold *α*_*FWE*_.

The formulation above clearly shows that the choice of the smoothing kernel has a critical impact on cluster inference. Spatial smoothing is considered as an essential pre-processing step to better comply with the RFT-assumptions (e.g. Hayasaka and Nichols, 2003). In Figure 1 the FWE corrected *p*-values are presented for a group analysis of 10 subjects over a hypothetical range of cluster sizes. In this analysis the data are smoothed with a kernel of 4 mm or 6 mm width. The derived smoothness (effective smoothness) then serves as the input for the formulas above.

Throughout this study, we consistently use a cluster-forming threshold that satisfies *P* (*T* ≥ *T*_*u*1_) = 0.001 as such threshold has better inferential properties (Hayasaka and Nichols, 2003; Eklund et al., 2016; Woo et al., 2014).

**Fig. 1.**
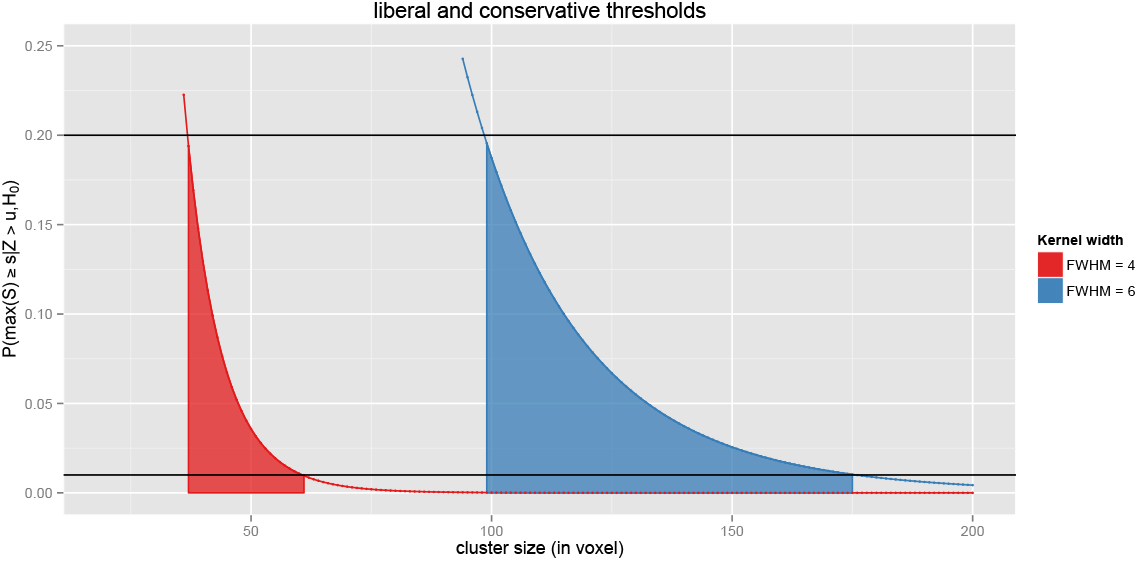
FWE corrected *p*-values in function of cluster size where the effective smoothness is based on an analysis of 10 subjects for an applied isotropic Gaussian smoothness kernel of 4mm and 6mm FWHM. The cluster-forming threshold satisfies *P* (*T* ≥ *T*_*u*1_) = 0.001. The filled area is the *deliberation* zone between the *liberal α*_*FWE*_ = 0.2 and *conservative α*_*FWE*_ = 0.01 threshold.

#### 2.2.2 Re-selection rate

For any given thresholding method, we define the re-selection rate (*rr*) at voxel level in a bootstrap resampling context. From a dataset at hand we draw *K* bootstrap samples. For each of the bootstrap samples, we re-run the original analysis that selects clusters. Next, we create a binary map in which the voxels belonging to a selected cluster are set to 1. Voxels that do not belong to such clusters are set to 0. In the final step we sum up the *K* binary maps (with a value *k* : 1*, …, K* per voxel) and divide these by *K*, resulting in a proportion for each voxel that indicates the re-selection rate in that voxel over bootstrap samples. The average re-selection rate per cluster is obtained by averaging over all voxels within the cluster.

#### 2.2.3 Procedure

We propose to set a *conservative* threshold at *α*_*FWE*_ = 0.01, and a *liberal* threshold at *α*_*FWE*_ = 0.2 for FWE corrected cluster *p*-values as defined in Equation 5. Clusters that survive the conservative threshold are selected. Clusters that do not survive the liberal threshold are not further considered. If a cluster only survives the liberal threshold, we let the cluster averaged reselection rate be the determining factor. For the clusters that lie in the deliberation zone between the two thresholds (see also Figure 1) are only selected if the average re-selection rate is larger than *π*_*thr*_, i.e. on average the voxels are selected in *π*_*thr*_ of the bootstrap samples. The heuristic is outlined below:

1. cluster survives the conservative threshold: **select the cluster**
2. cluster does not survive the liberal threshold: **do not select the cluster**
3. cluster only survives the liberal threshold: **select the cluster if average re-selection rate exceeds** *π*_*thr*_, with *π*_*thr*_ a pre-defined threshold (e.g. 2/3).

## 3 Evaluation of the procedure

In line with recent work on the evaluation of inferential fMRI procedures (Eklund et al., 2016, 2018; Nichols et al., 2017), we use real null and activation data. In all scenarios, stability was assessed on either *K* = 1000 or *K* = 500 bootstrap samples for respectively the null data and the activation data.

### 3.1 Operating characteristics

Using 2 real datasets, we evaluate if our method a) acceptably controls the amount of false positive findings and b) how the activation patterns are different from alternative inference strategies. While the first is assessed on fMRI data which mainly consist of noise (w.r.t. the imposed experiment design matrix), the latter is assessed on real activation fMRI data.

#### resting-state data

We assess the empirical FWE rate using resting-state data from the Cambridge subset (N=198) of the 1000 Functional Connectomes project. These data are obtained through www.nitrc.org. As no activation is present in these images, false positives are created using a random design matrix ***X*** (Eklund et al., 2016) for the determination of the *β* and the variance images that are propagated to the second level analysis. Pre-processing of this data is done similarly as in Eklund et al. (2016) but allowing for two different smoothing kernels: 4 and 6 mm wide FWHM (respectively 2 and 3 voxels width). The FWE is assessed for a group analysis with random subsamples of *n* = 10 or *n* = 20. Clusterwise FWE corrected *p*-values are obtained using FSL (Jenkinson et al., 2012). We assess the performance of the proposed procedure for the following parameter configurations: *π*_*thr*_ = (0.6, 2/3, 0.7, 0.8, 0.9) and thresholds pairs (*α*_*con*_ = 0.001, *α*_*lib*_ = 0.05); (*α*_*con*_ = 0.005,*α*_*lib*_ = 0.1); (*α*_*con*_ = 0.01,*α*_*lib*_ = 0.2) and (*α*_*con*_ = 0.05, *α*_*lib*_ = 0.4). For each parameter configuration, 1000 random samples are created to assess the average empirical FWE.

#### activation data

For the evaluation of our method, we use the 80 independent subjects package of the HCP (Van Essen et al., 2012) in a group analysis with randomly sampled samples of *n* = 10 or *n* = 20. We focus on 1 specific contrast of the emotion task, i.e. contrast 3 which compares faces versus shapes. The FWE-corrected *p*-values and the formation of the clusters are based on the FSL implemented command line tools (Jenkinson et al., 2012). We compare 4 thresholding methods: 1) a conservative 0.01 FWE-corrected threshold applied on the *p*-values derived from Equation 5 (*α*_*con*_ = 0.01); 2) the liberal 0.2 FWE-corrected threshold applied on the *p*-values derived from Equation 5 (*α*_*lib*_ = 0.2); 3) our above outlined balanced procedure with a liberal and conservative threshold, resp. *α*_*lib*_ = 0.2 and *α*_*con*_ = 0.01 (*α*_*lib*_&*π*_*trh*_ = 2/3) applied on the *p*-values derived from Equation 5; and 4) a procedure for which only clusters are selected that survive the liberal *α*_*FWE*_ = 0.2 and have an average re-selection rate that exceeds 2/3 (*π*_*trh*_ = 2/3). We add the fourth scenario to demonstrate how stability can also have a more prominent role in an inference strategy while ensuring that the FWE remains below a pre-specified level, which is nevertheless higher than a typical 5% level.

The following four thresholding methods are juxtaposed on 4 criteria: 1) the number of selected voxels; 2) the number of selected clusters; 3) the overlap between the mutually independent samples; and 4) the number of overlapping voxels between the mutually independent samples. The number of selected voxels/clusters is based on the respective thresholded test image from all 1000 samples.

The overlap between mutually independent samples is determined using the Jaccard Index (Jaccard, 1901):

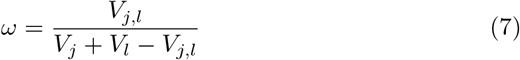

With 0 ≤ *ω*_*j,l*_ ≤ 1, and *V*_*j,l*_ the number of voxels in the union both images *j* and *l*. *V*_*j*_ and *V*_*l*_ denote respectively the total amount of active voxels in image *j* & *l*. *ω* thus denotes the ratio of the total amount of voxels that are selected from both test images versus the amount of selected voxels in each of the images.

The evaluation is based on 100 mutually exclusive samples of *n* = 10 and *n* = 20. This means that no identical images are in the two samples drawn from the 80 independent subjects package of the HCP (Van Essen et al., 2012) that are contrasted for the activation pattern.

### 3.2 Illustration

The re-selection rate can be easily represented graphically by the method proposed by Allen et al. (2012). By adding the re-selection as the level of transparency to a SPM heat map, a more voxel-based interpretation is added to the difficult interpretation of cluster-based inference plots. Note that this illustration is only based on one sample from the HPC data with *n* = 10.

## 4 Results

### 4.1 null data

The results for the pairwise comparisons of the mutually exclusive samples can be found in Table 2 for smoothing kernels with an FWHM of 4 or 6mm and *n* = 10 or 20.

**Table 2.** Family-wise Error Rates. *α*_*con*_.: conservative FWE-corrected threshold; *α*_*lib*_.: lib-eral FWE-corrected threshold.

| $(\alpha_{con.}; \alpha_{lib.})$ | no $\pi_{thr}$ | $\pi_{thr} = 0.6$ | $\pi_{thr} = 2/3$ | $\pi_{thr} = 0.7$ | $\pi_{thr} = 0.8$ | $\pi_{thr} = 0.9$ |
| --- | --- | --- | --- | --- | --- | --- |
| FWHM=4mm, n=10 |  |  |  |  |  |  |
| (0.001; 0.05) | 0.00000 | 0.00801 | 0.00501 | 0.00200 | 0.00000 | 0.00000 |
| (0.005; 0.1) | 0.00300 | 0.01301 | 0.00901 | 0.00501 | 0.00300 | 0.00300 |
| (0.01 ; 0.2) | 0.00300 | 0.01802 | 0.01201 | 0.00901 | 0.00300 | 0.00300 |
| (0.05 ; 0.4) | 0.01502 | 0.03804 | 0.02603 | 0.02302 | 0.01602 | 0.01502 |
| FWHM=4mm, n=20 |  |  |  |  |  |  |
| (0.001; 0.05) | 0.00402 | 0.01811 | 0.00704 | 0.00604 | 0.00402 | 0.00402 |
| (0.005; 0.1) | 0.01207 | 0.03018 | 0.01811 | 0.01610 | 0.01207 | 0.01207 |
| (0.01 ; 0.2) | 0.01610 | 0.04125 | 0.02414 | 0.02113 | 0.01610 | 0.01610 |
| (0.05 ; 0.4) | 0.04628 | 0.06740 | 0.05231 | 0.05030 | 0.04628 | 0.04628 |
| FWHM=6mm, n=10 |  |  |  |  |  |  |
| (0.001; 0.05) | 0.00302 | 0.01512 | 0.01411 | 0.01109 | 0.00302 | 0.00302 |
| (0.005; 0.1) | 0.00504 | 0.02319 | 0.01815 | 0.01411 | 0.00605 | 0.00504 |
| (0.01 ; 0.2) | 0.00706 | 0.04234 | 0.02823 | 0.02419 | 0.00907 | 0.00706 |
| (0.05 ; 0.4) | 0.01915 | 0.07560 | 0.05040 | 0.04133 | 0.02117 | 0.01915 |
| FWHM=6mm, n=20 |  |  |  |  |  |  |
| (0.001; 0.05) | 0.00800 | 0.03400 | 0.01600 | 0.00800 | 0.00800 | 0.00800 |
| (0.005; 0.1) | 0.01800 | 0.04700 | 0.02600 | 0.01800 | 0.01800 | 0.01800 |
| (0.01 ; 0.2) | 0.02300 | 0.05700 | 0.03100 | 0.02400 | 0.02300 | 0.02300 |
| (0.05 ; 0.4) | 0.05900 | 0.08200 | 0.06400 | 0.06100 | 0.05900 | 0.05900 |

For a low degree of smoothing, we find that with smaller sample sizes closer to nominal conservative FWE rates are obtained. Furthermore, we find that increasing the *π_*thr*_* results in situations where the threshold is too high, and results coincide with the situation where no threshold is set, i.e. the conservative threshold. Indeed, when *π*_*thr*_ = 2/3 we find a large decrease in FWE compared to *π*_*thr*_ = 0.6, as from *π*_*thr*_ = 0.8 the situation is highly similar to not using the additional information of the re-selection rate. When using a larger smoothing kernel with a FWHM of 6 mm, result show a similar pattern.

### 4.2 activation data

#### 4.2.1 *Sample with n* = 10

The results for the pairwise comparisons of the mutually exclusive samples with *n* = 10 can be found in Tables 3 for a smoothing kernel of 4 and 6mm. In general, we find that the liberal FWE threshold of *α* = 0.2 results in the highest number of selected clusters and voxels, while the conservative threshold of *α* = 0.01 results in the smallest number of selected clusters and voxels. Obviously, we find that for the strategy that uses two thresholds, the number of selected clusters (and thus voxels) lies between these numbers of the two thresholds. Compared to FWE control on cluster level with *α* = 0.01, we select more clusters but not at the cost of a substantial decline in the average overlap *ω*. On average we find about one half cluster extra if the data was smoothed with a 4mm width kernel while the difference is much smaller when a smoothing kernel of 6 mm was used.

**Table 3.**
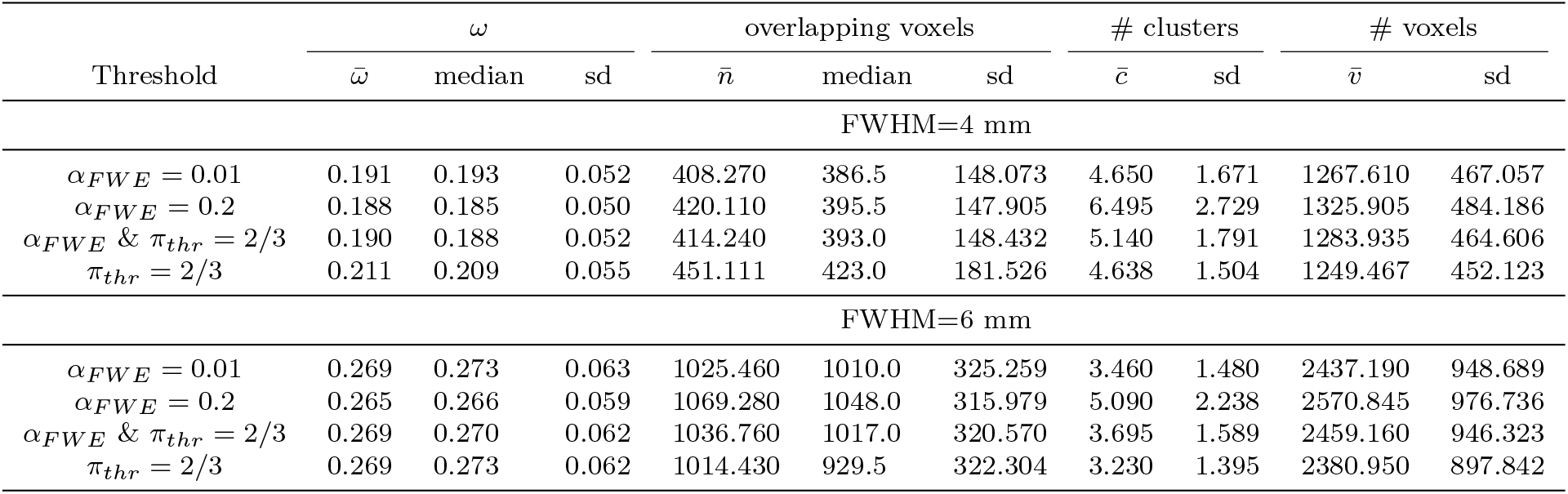
The average, median and standard deviation (sd) for *ω*, the number over overlapping voxels, the number of thresholded clusters and the number of thresholded voxels based on 100 mutually exclusive samples (*n* = 10) of the emotion task from the HPC data for an applied smoothing kernel of respectively 4 and 6 mm. *α*_*FWE*_ = 0.01: thresh-olded clusters by *α*_*FWE*_ = 0.01; *α*_*FWE*_ = 0.2: thresholded clusters by *α*_*FWE*_ = 0.2; *α*_*FWE*_ & *π*_thr_ > 2/3: thresholded clusters by either 1) *α*_*FWE*_ = 0.01 or 2)*α*_*FWE*_ = 0.2 and *π*_*thr*_ = 2/3; *π*_*thr*_ = 2/3: thresholded clusters by *α*_*FWE*_ = 0.2 and *π*_*thr*_ = 2/3.

| Threshold | $\omega$ | | | overlapping voxels | | | # clusters | | # voxels | |
| --- | --- | --- | --- | --- | --- | --- | --- | --- | --- | --- |
| | $\bar{\omega}$ | median | sd | $\bar{n}$ | median | sd | $\bar{c}$ | sd | $\bar{v}$ | sd |
| FWHM=4 mm |  |  |  |  |  |  |  |  |  |  |
| $\alpha_{FWE} = 0.01$ | 0.191 | 0.193 | 0.052 | 408.270 | 386.5 | 148.073 | 4.650 | 1.671 | 1267.610 | 467.057 |
| $\alpha_{FWE} = 0.2$ | 0.188 | 0.185 | 0.050 | 420.110 | 395.5 | 147.905 | 6.495 | 2.729 | 1325.905 | 484.186 |
| $\alpha_{FWE} \& \pi_{thr} = 2/3$ | 0.190 | 0.188 | 0.052 | 414.240 | 393.0 | 148.432 | 5.140 | 1.791 | 1283.935 | 464.606 |
| $\pi_{thr} = 2/3$ | 0.211 | 0.209 | 0.055 | 451.111 | 423.0 | 181.526 | 4.638 | 1.504 | 1249.467 | 452.123 |
| FWHM=6 mm |  |  |  |  |  |  |  |  |  |  |
| $\alpha_{FWE} = 0.01$ | 0.269 | 0.273 | 0.063 | 1025.460 | 1010.0 | 325.259 | 3.460 | 1.480 | 2437.190 | 948.689 |
| $\alpha_{FWE} = 0.2$ | 0.265 | 0.266 | 0.059 | 1069.280 | 1048.0 | 315.979 | 5.090 | 2.238 | 2570.845 | 976.736 |
| $\alpha_{FWE} \& \pi_{thr} = 2/3$ | 0.269 | 0.270 | 0.062 | 1036.760 | 1017.0 | 320.570 | 3.695 | 1.589 | 2459.160 | 946.323 |
| $\pi_{thr} = 2/3$ | 0.269 | 0.273 | 0.062 | 1014.430 | 929.5 | 322.304 | 3.230 | 1.395 | 2380.950 | 897.842 |

#### 4.2.2 *Sample with n* = 20

The results for the pairwise comparisons of the mutually exclusive samples with *n* = 20 can be found in Tables 4 for a smoothing kernel of 4 and 6mm. In general we find a similar pattern as with a sample size of *n* = 10. We note however that with a smoothing kernel of 6 mm the average increase in selected voxels is about 100 when using the balanced criterion instead of an FWE control with *α* = 0.001. In contrast to the sample size of *n* = 10, our procedure results on average in about 1.5 extra clusters, with a total additional number of voxels ranging up to on average about 100 voxels extra if a smoothing kernel width of 6 mm is used.

**Table 4.**
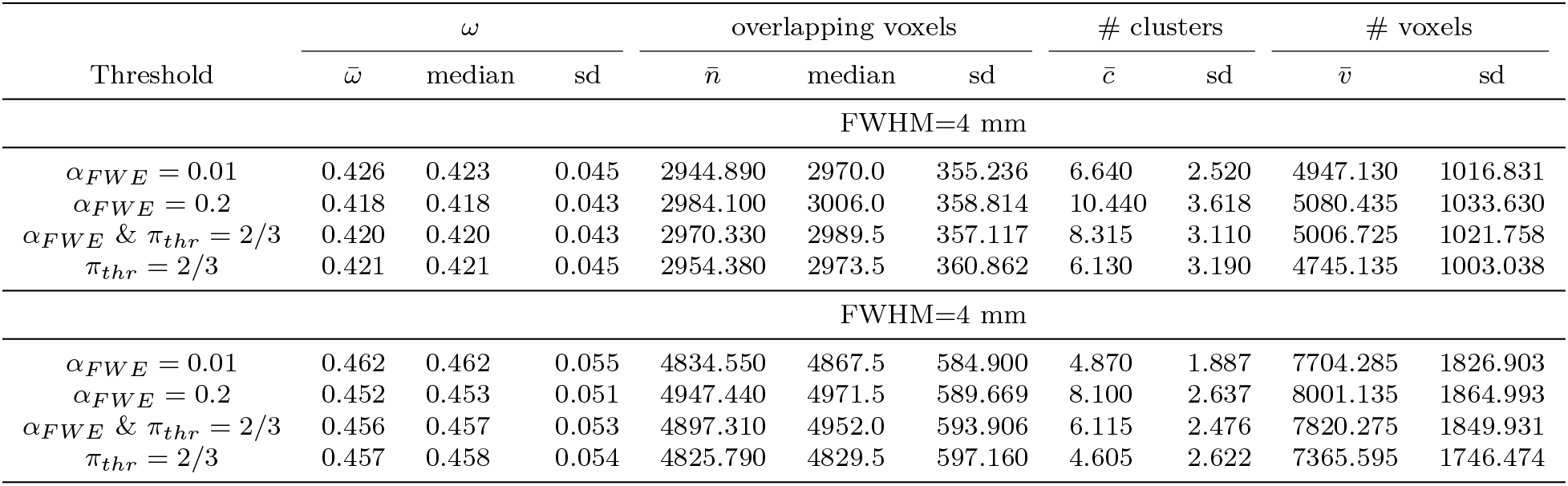
The average, median and standard deviation (sd) for *ω*, the number over overlapping voxels, the number of thresholded clusters and the number of thresholded voxels based on 100 mutually exclusive samples (*n* = 20) of the emotion task from the HPC data for an applied smoothing kernel of respectively 4 and 6 mm. *α*_*FWE*_ = 0.01: thresh-olded clusters by *α*_*FWE*_ = 0.01; *α*_*FWE*_ = 0.2: thresholded clusters by *α*_*FWE*_ = 0.2; *α*_*FWE*_ & *π*_*thr*_ > 2/3: thresholded clusters by either 1) *α*_*FWE*_ = 0.01 or 2)*α*_*FWE*_ = 0.2 and *π_thr_* = 2/3; *π*_*thr*_ = 2/3: thresholded clusters by *α*_*FWE*_ = 0.2 and *π*_*thr*_ = 2/3.

| Threshold | $\omega$ | | | overlapping voxels | | | # clusters | | # voxels | |
| --- | --- | --- | --- | --- | --- | --- | --- | --- | --- | --- |
| | $\bar{\omega}$ | median | sd | $\bar{n}$ | median | sd | $\bar{c}$ | sd | $\bar{v}$ | sd |
| FWHM=4 mm |  |  |  |  |  |  |  |  |  |  |
| $\alpha_{FWE} = 0.01$ | 0.426 | 0.423 | 0.045 | 2944.890 | 2970.0 | 355.236 | 6.640 | 2.520 | 4947.130 | 1016.831 |
| $\alpha_{FWE} = 0.2$ | 0.418 | 0.418 | 0.043 | 2984.100 | 3006.0 | 358.814 | 10.440 | 3.618 | 5080.435 | 1033.630 |
| $\alpha_{FWE} \& \pi_{thr} = 2/3$ | 0.420 | 0.420 | 0.043 | 2970.330 | 2989.5 | 357.117 | 8.315 | 3.110 | 5006.725 | 1021.758 |
| $\pi_{thr} = 2/3$ | 0.421 | 0.421 | 0.045 | 2954.380 | 2973.5 | 360.862 | 6.130 | 3.190 | 4745.135 | 1003.038 |
| FWHM=6 mm |  |  |  |  |  |  |  |  |  |  |
| $\alpha_{FWE} = 0.01$ | 0.462 | 0.462 | 0.055 | 4834.550 | 4867.5 | 584.900 | 4.870 | 1.887 | 7704.285 | 1826.903 |
| $\alpha_{FWE} = 0.2$ | 0.452 | 0.453 | 0.051 | 4947.440 | 4971.5 | 589.669 | 8.100 | 2.637 | 8001.135 | 1864.993 |
| $\alpha_{FWE} \& \pi_{thr} = 2/3$ | 0.456 | 0.457 | 0.053 | 4897.310 | 4952.0 | 593.906 | 6.115 | 2.476 | 7820.275 | 1849.931 |
| $\pi_{thr} = 2/3$ | 0.457 | 0.458 | 0.054 | 4825.790 | 4829.5 | 597.160 | 4.605 | 2.622 | 7365.595 | 1746.474 |

From Tables 3–4 it is interesting to note that when *π*_*thr*_ = 2/3 is used alone in combination with the liberal threshold a smaller amount of clusters is selected. However, a good overlap between the samples is found.

#### 4.2.3 Illustration

The method and implementation of Allen et al. (2012) enables high dimensional visualisation of data properties. We use this principle to incorporate data analytical stability into classical fMRI plots using heat maps on brain slices. On Figure 2 we distinguish 3 different layers: 1) the classical heat map on the color scale with a Z-statistic; 2) the data analytical stability displayed as the transparency of the colors and 3) the contours indicate the selected clusters. We demonstrate the principle for the analysis for one sample with *n* = 10, using a smoothing kernel of respectively 4 mm and 6 mm width.

**Fig. 2.**
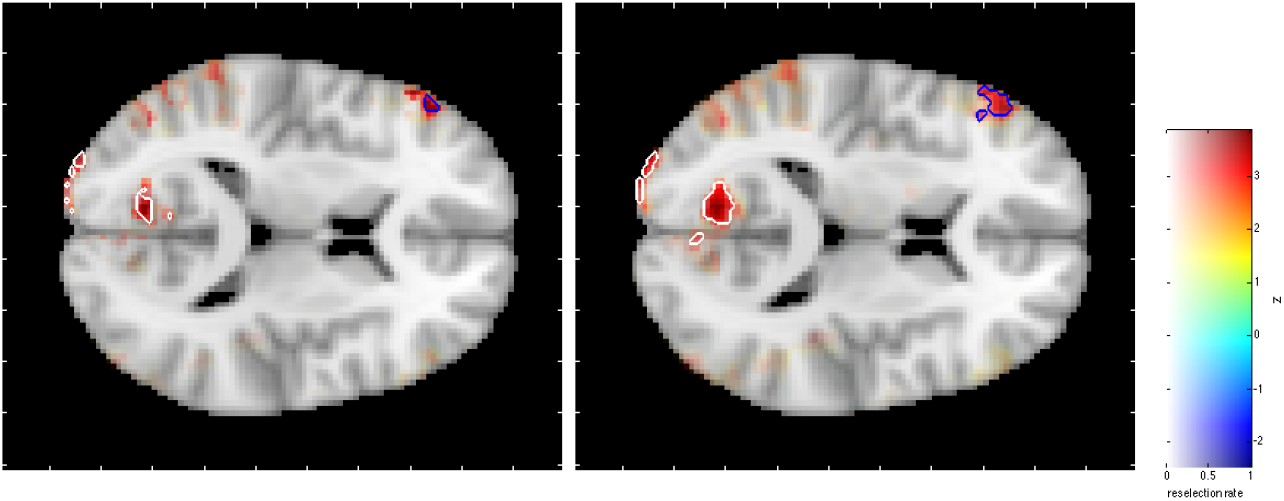
Activation for contrast 3 in the HCP data. The white contours indicate clusters that are selected based on cluster-based inference with *α*_*FWE*_ = 0.01. The blue contour indicates a cluster that only survives the liberal threshold, but also satisfies *π*_*thr*_ = 2/3. The colors correspond with the value of the *Z* statistic, more transparency indicates a lower re-selection rate.

As two analyses only differ in the amount of smoothing applied, the results in Tables 5 and 6 highly resemble each other, which is also clear in the left and the middle panel from Figure 2. Based on the results in Table 5, next to selection of clusters 5.1-5.3, we additionally select cluster 5.5, which has a *p* = 0.079 but an average re-selection rate of 0.827. It is furthermore remarkable that with a smoothing kernel of 6 mm, a similar pattern occurs, in which cluster 6.4 is added to the selection. Note however the substantial difference in size due to the choice of kernel width.

**Table 5.** Results from 1 FSL-based cluster analysis for *n* = 10 with a smoothing kernel of 4 and 6 mm. With *α*_*con*_. = 0.01 and *α*_*lib*_ = 0.2.

| | ID | size | $maxZ$ | FWE $p$ | X | Y | Z | average re-selection rate |
| --- | --- | --- | --- | --- | --- | --- | --- | --- |
| survive $\alpha_{con.}$ | 5.1 | 790 | 4.750 | $< 0.0001$ | 31 | 17 | 33 | 0.724 |
| | 5.2 | 375 | 5.330 | $< 0.0001$ | 66 | 21 | 30 | 0.774 |
|  | 5.3 | 64 | 4.800 | 0.001 | 64 | 36 | 27 | 0.836 |
| survive $\alpha_{lib.}$ | 5.4 | 31 | 4.130 | 0.058 | 36 | 63 | 28 | 0.510 |
|  | **5.5** | 29 | 4.010 | **0.079** | 20 | 83 | 41 | **0.827** |
|  | 5.6 | 26 | 3.740 | 0.127 | 56 | 29 | 39 | 0.365 |

**Table 6.**
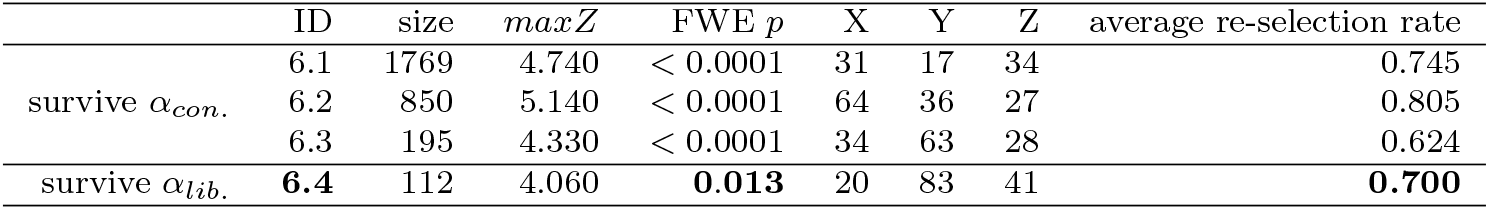
Results from 1 FSL-based cluster analysis for *n* = 10 with a smoothing kernel of 6 mm. With *α*_*con*_. = 0.01 and *α*_*lib*_. = 0.2.

| | ID | size | $maxZ$ | FWE $p$ | X | Y | Z | average re-selection rate |
| --- | --- | --- | --- | --- | --- | --- | --- | --- |
| survive $\alpha_{con.}$ | 6.1 | 1769 | 4.740 | $< 0.0001$ | 31 | 17 | 34 | 0.745 |
| | 6.2 | 850 | 5.140 | $< 0.0001$ | 64 | 36 | 27 | 0.805 |
| | 6.3 | 195 | 4.330 | $< 0.0001$ | 34 | 63 | 28 | 0.624 |
| survive $\alpha_{lib.}$ | **6.4** | 112 | 4.060 | **0.013** | 20 | 83 | 41 | **0.700** |

The visualization in Figure 2 also allows to explore the data analytical stability in specific regions. For the selected clusters, we see that those at the borders are less stable, especially with a kernel width of 4 mm. Furthermore, Figure 2 allows to inspect the data analytical stability of voxels that do not exceed the the cluster-forming threshold. Based on these figures, we can seem both for 4 and 6 mm smoothing, relatively stable activation in the left temporal cortex.

## 5 Discussion

We presented a testing procedure for the selection of clusters that balances between sufficient control on the amount of false positives and sufficient ability to detect activation by considering stability of the activation. We retain clusters that either survive the conservative FWE threshold or survive the liberal FWE threshold and also have a high average re-selection rate. Our results show that this procedure is successful in the selection of more clusters and, consequently more voxels are declared active without substantial losses on the operating characteristics compared to similar thresholding methods in null, resting-state, data.

By setting a theoretical framework to define the two thresholds our proposal intrinsically takes into account the properties of the data (i.e. the size of the volume, the smoothness and the height of the first thresholds). Given the current controversies on the reliance on such theoretical framework and it dependence on data-characteristics (Eklund et al., 2016, 2018), we add the rationale for selecting relevant features in line with the original work of Meinshausen and Buühlmann (2010) on stability of feature extraction. Secondly, the addition of a voxel-wise metric within the cluster-based inference increases the interpret-ability of the voxels *in* a cluster. In two ways we find more information compared to classical testing frameworks. Within a selected cluster our procedure allows to further differentiate between within-cluster regions that are highly stable versus regions that not stable. Also, with the re-selection rates we can highlight regions that even do not survive liberal threshold but are relatively stable and would otherwise not have been picked up. It should be stressed that even with the current call to have more stringent control on the False Positives (Benjamin and Johnson, 2017), we would lack such advantages.

While the concept of data analytical stability has recently been introduced mainly for the evaluation of methodological choices in fMRI data analysis (Roels et al., 2015, 2016), it has previously been used to improve corrections for multiple testing (e.g. Gordon et al., 2009) also in fMRI (Durnez et al., 2014b). While computationally intensive, the implementation of data analytical stability in the voxel-wise single-subject analysis of fMRI was successful in the improvements of stability of the FDR correction for multiple testing (Durnez et al., 2014b). Although our method is also computationally intensive, it is less computationally intensive compared to first-level modeling because of several reasons: the tacit assumption that all first-level modeling is correct, less observations (time points versus study participants) and typically independent observations.

As the inferential validity of cluster-based inference might not be guaranteed for low first thresholds or too low smoothing (Eklund et al., 2016; Hayasaka and Nichols, 2003), we insure this validity by setting a high cluster-forming threshold (*α* = 0.001) and incorporating the data analytical stability. Our method is however sufficiently flexible to use permutation-based cluster-forming thresholds. Moreover, the inclusion of the average re-selection of the voxels within a cluster provides a useful metric for the potential instability of the cluster-based thresholds in that with low intrinsic smoothing.

At last we stress that the proposed procedure is relatively immune to so-called *p*-hacking, the coarse practice to smuggle in significance of results that lie near a fixed threshold. We found good performance of our procedure with setting the cluster-forming threshold at 0.001, and imposing a the conservative threshold at 0.01 and the liberal threshold at 0.2 for the FWE corrected cluster *p*-values. In the light of a recent call to pre-register scientific studies (e.g. Nosek et al., 2018), and the required predetermined choices for all statistical thresholds, with this procedure we emphasize the importance of avoiding posthoc adjustments to these thresholds. Also by re-sampling the data to obtain the re-selection rates, we clearly choose for a more robust approach. At last, the use of auxiliary voxel-wise information in a cluster-based inference framework enriches the classical binary decision framework in determining active clusters, while controlling the amount of FP’s in a principled way.

## Conclusion

In this paper we have presented a procedure that successfully balances between sufficient control on the amount of false positives and sufficiently high power by including data analytical stability in cluster-based inference. The inclusion of data analytical stability additionally aids the interpretation of the results.

## Acknowledgements

The computational resources (Stevin Supercomputer Infrastructure) and services used in this work were provided by the VSC (Flemish Supercomputer Center), funded by Ghent University, the Hercules Foundation and the Flemish Government department EWI.

NITRC, NITRC-IR, and NITRC-CE have been funded in whole or in part with Federal funds from the the Department of Health and Human Services, National Institute of Biomedical Imaging and Bioengineering, the National Institute of Neurological Disorders and Stroke, under the following NIH grants: 1R43NS074540, 2R44NS074540, and 1U24EB023398 and previously GSA Contract No. GS-00F-0034P, Order Number HHSN268200100090U

## References

Allen, E. a., Erhardt, E. B., and Calhoun, V. D. (2012). Data Visualization in the Neurosciences: Overcoming the Curse of Dimensionality. Neuron, 74(4):603–608.

Beckmann, C. F., Jenkinson, M., and Smith, S. M. (2003). General multilevel linear modeling for group analysis in FMRI. NeuroImage, 20(2):1052–63.

Benjamin, D. J. B. J. J. M. N. B. A. W. E. . . and Johnson, V. (2017). Redefine statistical significance. PsyArXiv.

Benjamini, Y. and Hochberg, Y. (1995). Controlling the False Discovery Rate: A Practical and Powerful Approach to Multiple Testing.

Brett, M., Penny, W., and Kiebel, S. (2007). Parametric procedures. In Friston, K. J., Ashburner, J., Kiebel, S., Nichols, T., and Penny, W., editors, Statistical Parametric Mapping: The Analysis of Functional Brain Images, chapter 8. Elsevier Ltd./Academic Press.

Button, K. S., Ioannidis, J. P. a., Mokrysz, C., Nosek, B. a., Flint, J., Robinson, E. S. J., and Munafo, M. R. (2013). Power failure: why small sample size undermines the reliability of neuroscience. Nature reviews. Neuroscience, 14(5):365–76.

Carp, J. (2012). The secret lives of experiments: Methods reporting in the fMRI literature. NeuroImage, 63(1):289–300.

Chumbley, J., Worsley, K., Flandin, G., and Friston, K. (2010). Topological FDR for neuroimaging. NeuroImage, 49(4):3057–64.

Chumbley, J. R. and Friston, K. J. (2009). False discovery rate revisited: FDR and topological inference using Gaussian random fields. NeuroImage, 44(1):62–70.

Cochrane, D. and Orcutt, G. (1949). Application of least squares regression to relationships containing auto-correlated error terms. Journal of the American Statistical Association, 44(245):32–61.

Durnez, J., Moerkerke, B., and Nichols, T. E. (2014a). Post-hoc power estimation for topological inference in fMRI. NeuroImage, 84:45–64.

Durnez, J., Roels, S., and Moerkerke, B. (2014b). Multiple testing in fmri: a case study on the balance between sensitivity, specificity and stability. Biometrical Journal, 56(4).

Eklund, A., Nichols, T. E., and Knutsson, H. (2016). Cluster failure: Why fMRI inferences for spatial extent have inflated false-positive rates. Proceedings of the National Academy of Sciences, page 201602413.

Eklund, A., Nichols, T. E., and Knutsson, H. (2018). Cluster failure revisited: Impact of first level design and physiological noise on cluster false positive rates. Human Brain Mapping, 40(11).

Forman, S. D., Cohen, J. D., Fitzgerald, M., Eddy, W. F., Mintun, M. A., and Noll, D. C. (1995). Improved assessment of significant activation in functional magnetic resonance imaging (fmri): Use of a cluster-size threshold. Magnetic Resonance in Medicine, 33(5):636–647.

Friston, K. J., Holmes, a., Poline, J. B., Price, C. J., and Frith, C. D. (1996). Detecting activations in PET and fMRI: levels of inference and power. Neu-roImage, 4(3 Pt 1):223–235.

Gordon, A., Chen, L., Glazko, G., and Yakovlev, A. (2009). Balancing Type One and Two Errors in Multiple Testing for Differential Expression of Genes. Computational statistics & data analysis, 53(5):1622–1629.

Gordon, A., Glazko, G., Qiu, X., and Yakovlev, A. (2007). Control of the mean number of false discoveries, Bonferroni and stability of multiple testing. The Annals of Applied Statistics, 1(1):179–190.

Hayasaka, S. and Nichols, T. E. (2003). Validating cluster size inference: random field and permutation methods. NeuroImage, 20(4):2343–2356.

Henson, R. and Friston, K. J. (2007). Convolution models for fmri. In Friston, K., Ashburner, J., Kiebel, S., Nichols, T., and Penny, W., editors, Statistical Parametric Mapping: The Analysis of Functional Brain Images, pages 193–210. Elsevier Ltd./Academic Press.

Jaccard, P. (1901). Distribution florale dans une portion des alpes et du jura. Bulletin de la Société Vaudoise des Sciences Naturelles, 37:547–579.

Jenkinson, M., Beckmann, C. F., Behrens, T. E. J., Woolrich, M. W., and Smith, S. M. (2012). Fsl. NeuroImage, 62(2):782–90.

Kiebel, S. and Holmes, A. P. (2007). The general linear model. In Friston, K. J., Ashburner, J., Kiebel, S., Nichols, T., and Penny, W., editors, Statistical Parametric Mapping: The Analysis of Functional Brain Images, chapter 8. Elsevier Ltd./Academic Press.

Lindquist, M. A. (2008). The Statistical Analysis of fMRI Data. Statistical Science, 23(4):439–464.

Meinshausen, N. and Buühlmann, P. (2010). Stability selection. Journal of the Royal Statistical Society: Series B (Statistical Methodology), 72(4):417–473.

Mumford, J. A. (2012). A power calculation guide for fMRI studies. Social cognitive and affective neuroscience, 7(6):738–42.

Mumford, J. A. and Nichols, T. E. (2006). Modeling and inference of multi-subject fMRI data. IEEE Engineering in Medicine and Biology Magazine, 25(2):42–51.

Mumford, J. A. and Nichols, T. E. (2009). Simple group fMRI modeling and inference. NeuroImage, 47(4):1469–75.

Nichols, T. E. (2012). Multiple testing corrections, nonparametric methods, and random field theory. NeuroImage, 62(2):811–5.

Nichols, T. E., Eklund, A., and Knutsson, H. (2017). A defense of using resting-state fmri as null data for estimating false positive rates. Cognitive Neuroscience, 8(3):144–149. PMID: 28140785.

Nichols, T. E. and Hayasaka, S. (2003). Controlling the familywise error rate in functional neuroimaging: a comparative review. Statistical methods in medical research, 12(5):419–46.

Nosek, B., Ebersole, C. R., DeHave, A. C., and Mellor, D. T. (2018). The preregistration revolution. Procedures of the National Academy of Sciences, 115(11).

Poldrack, R. A., Mumford, J. A., and Nichols, T. E. (2011). Handbook of Functional MRI Data Analysis. Cambridge University Press, New York.

Poline, J.-B. and Brett, M. (2012). The general linear model and fMRI: Does love last forever? NeuroImage, 62(2):871–880.

Qiu, X., Xiao, Y., Gordon, A., and Yakovlev, A. (2006). Assessing stability of gene selection in microarray data analysis. BMC Bioinformatics, 7.

Roels, S. P., Loeys, T., and Moerkerke, B. (2016). Evaluation of Second-Level Inference in fMRI Analysis. Computational Intelligence and Neuroscience, 2016(Article ID 1068434):22.

Roels, S. P., Moerkerke, B., and Loeys, T. (2015). Bootstrapping fmri data: Dealing with misspecification. Neuroinformatics, 13(3):337–352.

Van Essen, D. C., Ugurbil, K., Auerbach, E., Barch, D., Behrens, T. E. J., Bucholz, R., Chang, a., Chen, L., Corbetta, M., Curtiss, S. W., Della Penna, S., Feinberg, D., Glasser, M. F., Harel, N., Heath, a. C., Larson-Prior, L., Marcus, D., Michalareas, G., Moeller, S., Oostenveld, R., Petersen, S. E., Prior, F., Schlaggar, B. L., Smith, S. M., Snyder, a. Z., Xu, J., and Yacoub, E. (2012). The Human Connectome Project: A data acquisition perspective. NeuroImage, 62(4):2222–2231.

Woo, C.-W., Krishnan, A., and Wager, T. D. (2014). Cluster-extent based thresholding in fMRI analyses: pitfalls and recommendations. NeuroImage, 91:412–9.

Worsley, K. J. (2007). Random field theory. In Friston, K., Ashburner, J., Kiebel, S., Nichols, T., and Penny, W., editors, Statistical Parametric Mapping: The Analysis of Functional Brain Images, pages 232–236. Elsevier Ltd./Academic Press.

Worsley, K. J., Liao, C. H., Aston, J., Petre, V., Duncan, G. H., Morales, F., and Evans, A. C. (2002). A general statistical analysis for fMRI data. NeuroImage, 15(1):1–15.

